# A novel behavioral paradigm to model hypermemory-like features in rats

**DOI:** 10.64898/2026.10.08.757535

**Authors:** Elvira Conforti, Giulia Chiacchierini, Alberto Marchetti, Jacopo Maria Lisiola, Valerio Santangelo, Patrizia Campolongo

## Abstract

Episodic memory is essential to everyday life and personal identity. However, its dysfunctions can profoundly affect mental health, and translational methods for investigating them in animals remain limited. Most rodent paradigms focus on memory deficits, whereas few tools model abnormally enhanced and persistent memories. Here, we developed a behavioral paradigm to induce hypermemory-like features in adult male and female rats using a version of object recognition (OR) and object location (OL) tasks specifically modified to mimic the rumination-like component of hypermemory. During the odor-context association training, rats explored two identical objects in the presence of a cinnamon scent. Over the following 6 h, they received 12 brief re-exposures in their home cage to either the cinnamon odor or an odor-free stimulus. By repeatedly reactivating the memory trace, these re-exposures promoted its hyperconsolidation without re-exposing animals to the original context or objects. Twenty-four hours later, memory was assessed under odor-free conditions. Odor re-exposure selectively enhanced OL memory in rats of both sexes, whereas OR memory remained unaffected. The procedure did not alter total object exploration or locomotor activity. Enhanced spatial memory was also accompanied by increased c-Fos expression in the ventral hippocampus in both sexes. Our protocol recapitulates key features observed in individuals with hyperfunctional episodic memory: continuous memory reactivation and rumination which may contribute to the excessive strengthening and persistence of specific memories. This simple and reproducible paradigm provides a novel translational tool for investigating the behavioral and neurobiological mechanisms underlying memory hyperconsolidation and maladaptive memory persistence.

## 1. Introduction

Episodic memory, a subtype of declarative memory, enables individuals to mentally reconstruct past personal experiences that occurred at specific times and places (Tulving, 1983; Squire and Zola, 1996; Tulving, 2002). This form of memory relies on the coordinated activity of distributed brain networks, particularly involving the hippocampus and prefrontal cortex, to support encoding, retrieval, and contextual processing (Rugg and Vilberg, 2013; Sugar and Moser, 2019; Danieli et al., 2023). Moreover, episodic memories do not remain unchanged over time; rather, they undergo reconsolidation and modification processes based on new experiences, which can strengthen, weaken, or integrate additional information into existing knowledge (Nadel et al., 2012; Danieli et al., 2023). Overall, the available literature highlights the complexity of this memory system.

While episodic memory has been extensively studied in humans, modeling its complexity in non-human animals remains a significant challenge. Importantly, existing animal models have largely focused on memory impairments (Khan et al., 2026), with relatively little attention given to conditions characterized by excessive or persistent memory (Santangelo et al., 2022). However, memory dysfunction in humans is not limited to memory loss; it can also manifest as an inability to suppress recollections of routine or futile memories. A striking example of this is a condition known as Highly Superior Autobiographical Memory (HSAM) (Parker et al., 2006; LePort et al., 2012; Santangelo et al., 2018; Santangelo et al., 2020; Santangelo et al., 2021), a rare condition in which individuals exhibit an exceptional ability to recall personal life events with high accuracy and detail. While HSAM is often described as a remarkable cognitive ability, it is also associated with increased rumination, obsessive-compulsive tendencies, and difficulty to disengage from past experiences, suggesting that enhanced memory retention may come at the cost of reduced cognitive flexibility (Parker et al., 2006; LePort et al., 2012; Santangelo et al., 2025; Pedale et al., 2026).

This aspect of memory persistence is particularly relevant in the context of neuropsychiatric disorders. Conditions such as post-traumatic stress disorder (PTSD), depression, obsessive-compulsive disorder (OCD), and anxiety disorders are frequently characterized by intrusive memories, rumination, and impaired forgetting (Morena et al., 2018; Park et al., 2024; Mattioni et al., 2025). Importantly, rumination is not restricted to traumatic memory, as observed in PTSD, but represents a transdiagnostic feature across different pathological conditions. These phenomena point to a failure of the mechanisms that normally support adaptive memory regulation, including the suppression or updating of irrelevant and maladaptive information (Donaldson and Hen, 2015; Schmitz et al., 2017).

Incorporating rumination-like, increased memory retention and resistance to updating components into animal models could therefore improve the alignment between preclinical and clinical research, facilitating the comparison between animal behavior and human cognitive-affective symptoms. Moreover, such a model may provide a useful platform to investigate the neurobiological mechanisms underlying maladaptive memory persistence and to test novel pharmacological or circuit-based interventions.

In this study we introduce a novel behavioral paradigm to model hypermemory-like features in rats. By using a forced rumination protocol, we aim to reproduce key aspects of enhanced memory retention and rumination observed in humans (Santangelo et al., 2018; Parente et al., 2026). This experimental design is built on the principle of forced hyperconsolidation: repeated post-training re-exposure to a learning-associated odor is designed to trigger iterative reactivation-consolidation events, progressively strengthening the memory trace. To this end, olfactory stimuli were selected as retrieval cues: their direct anatomical access to the entorhinal-hippocampal circuit confers exceptional efficacy in triggering contextual memory reactivation (Chapuis and Wilson, 2011; Aqrabawi and Kim, 2018), making repeated odor exposure a naturalistic tool for inducing iterative consolidation cycles. We hypothesized that repeated olfactory reactivation would selectively enhance hippocampus-dependent memory performance and engage ventral hippocampal circuitry, providing a rat model with face validity for the study of hypermemory-related phenomena.

## 2. Materials and Methods

### 2.1 Animals

Adult male and female Sprague-Dawley rats were used (250–300 g, ∼7-8 weeks, at the time of testing; Charles River Laboratories, Calco, Italy). Rats were kept in a temperature-controlled colony room (21 ± 1◦C; lights on from 07:00 AM to 7:00 PM) with food and water *ad libitum*. All the behavioral tests were performed during the light phase of the cycle. Animals were single housed starting from 3 days prior to behavioral tests until the end of experiments. We have previously shown that social isolation is necessary for the consolidation of long-lasting memories (Berardi et al., 2014). The animal protocol was approved by the Italian Ministry of Health. All procedures involving living animals were performed in compliance with the ARRIVE guidelines, the Directive 2010/63/EU of the European Parliament, and the D.L. 26/2014 of Italian Ministry of Health.

### 2.2 Behavioral procedures and equipment

During the 3 days preceding the beginning of the behavioral procedures, a cylindrical glass container equipped with a perforated metal lid (3,7 x 3,7 x 7,0 cm, 50 ml capacity) and filled with clean bedding was placed inside each home cage to habituate animals to this container used at a later time during the behavioral protocol. All the behavioral sessions were conducted in a gray Plexiglas box (40 × 40 × 40 cm) with the floor covered with a 2-cm layer of bedding, under dim light conditions (5 lux on the apparatus). Rats were individually handled for three days (1 min/day) and subsequently placed in the experimental arena for 3 min/day (**habituation**) to familiarize them to both the experimenter and the experimental context, thereby reducing procedure-related stress (Fig. 1). After each session, fecal boli were removed, the bedding was remixed, and the arena walls were cleaned with 70% ethanol (Campolongo et al., 2012; Tucci et al., 2014). Behavioral sessions were video-recorded and subsequently scored using ANY-maze software by blind experimenters to the experimental conditions.

**Fig 1.**
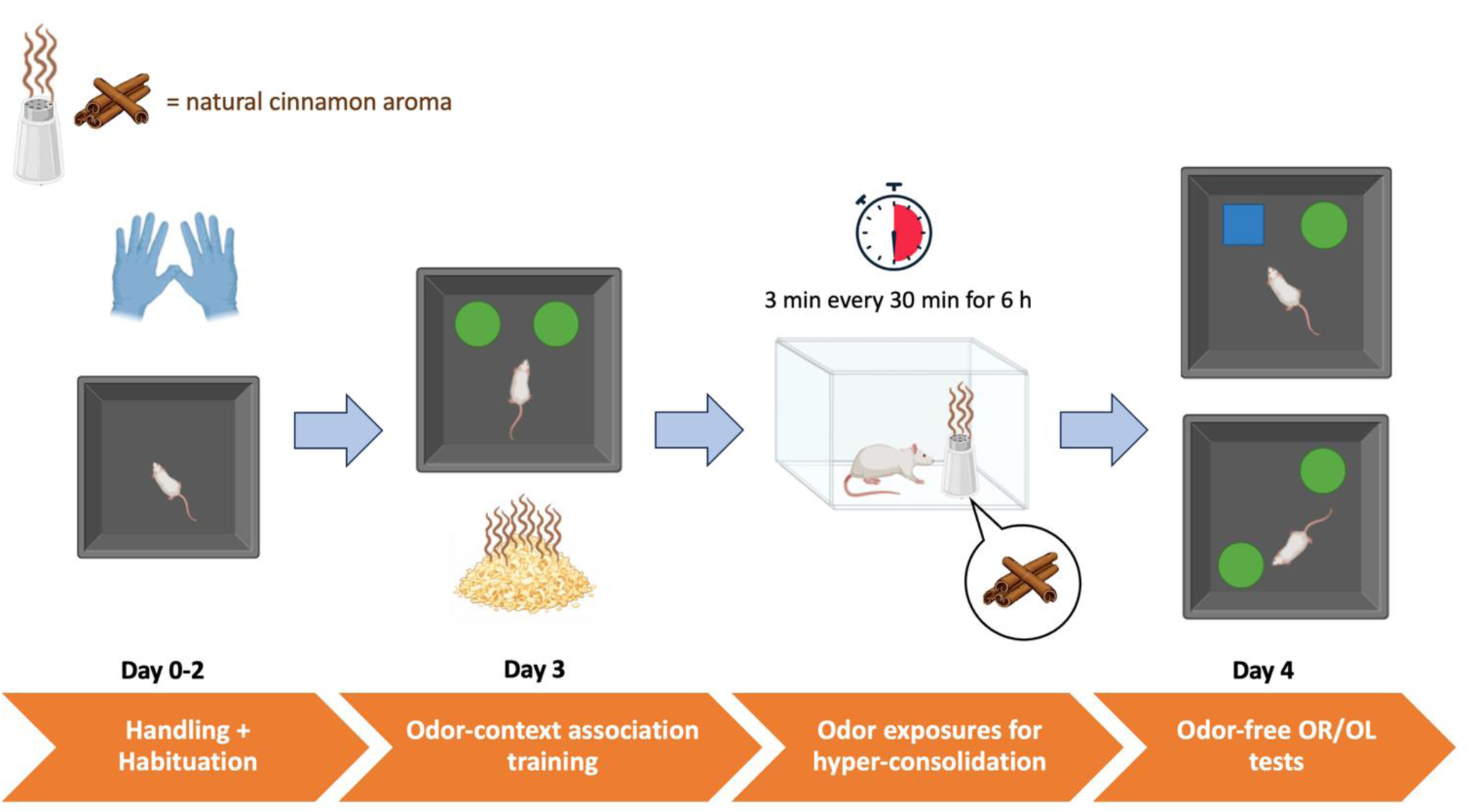
Schematic representation of the odor-based forced hyperconsolidation paradigm. Starting three days before the behavioral procedures, rats were handled for 1 min/day and habituated to the experimental arena for 3 min/day. During training, animals explored two identical objects for 3 min in an arena containing bedding impregnated with a 10% cinnamon solution. Over the following 6 h, rats in the odor re-exposure group (OD) were exposed in their home cage to cinnamon-scented bedding for 3 min every 30 min, whereas control animals (NO) were exposed to non-scented bedding. Twenty-four hours after training, memory was assessed using either the odor-free object recognition (OR) test, in which one object was replaced with a novel object, or the odor-free object location (OL) test, in which one object was moved to a novel position. No olfactory cues were present during testing. Memory performance was evaluated using the discrimination index.

#### Odor-context association training

Twenty-four hours after the last habituation session, each animal was individually placed in the center of the apparatus, with its back facing the objects, and allowed to freely explore two identical objects, A and B, positioned in opposite corners, for 3 min. Before the training session, the bedding covering the arena floor was impregnated with a 10% natural cinnamon flavor solution dissolved in mineral oil. The cinnamon solution was reapplied between sessions to maintain a stable odor intensity, inducing the animals to associate the odor to the context. Objects were not scented and were cleaned with 70% ethanol between animals. Object exploration was defined as the time during which the animal directed its nose toward the object from a distance of ≤ 1 cm (Campolongo et al., 2013; Tucci et al., 2014). At the end of each session, animals were returned to their home cages, and the arena was cleaned as described above. During the following 6 h, corresponding to the early post-training synaptic consolidation window in which newly encoded memory traces remain labile and susceptible to modulation (Dudai et al., 2004; Casagrande et al., 2018; Bellfy & Kwapis, 2020), at 30-min intervals, animals in the “odor re-exposure” group (OD) were exposed in their home cage to the glass container filled with bedding impregnated with the same cinnamon/mineral oil solution used during training. Each odor re-exposure lasted 3 minutes, and animals received in total 12 re-exposures. Control animals (“no odor”, NO) were exposed, at identical timings, to a container filled with non-scented bedding. After the final exposure session, all containers were removed.

#### Odor-free Object Recognition / Object Location tests

Twenty-four hours after training, animals underwent either the object recognition or object location test. Animals were reintroduced into the apparatus in the same orientation used during training and allowed to explore the objects for 3 min. In the object recognition paradigm, one familiar object, A, and one novel object , C, were presented in the same positions used during training. In the object location paradigm, both objects were familiar, but one object was displaced to a novel location while the other remained in its original position. During the test phase, neither the bedding nor the objects were treated with olfactory cues. Exploration time of the familiar object/object location (Told/Tnon-displaced) and the novel ones (Tnovel/Tdisplaced), and general exploratory activity were recorded. General exploratory activity was assessed by scoring the number of crossings and the frequency and duration of rearings and wall-rearings behaviors. Memory performance was quantified using the discrimination index: DI = (Tnovel − Told) / (Tnovel + Told) for object recognition, and DI = (Tdisplaced − Tnon-displaced) / (Tdisplaced + Tnon-displaced) for object location. Animals were excluded from the analysis if they showed health or welfare conditions incompatible with behavioral testing, if they failed to explore both objects during training or test (0 s of exploration on either object), or if their total object exploration time was <10s (**Fig. 1A**).

### 2.3 Immunohistochemistry

At the end of the experimental procedures, animals were euthanized with an overdose of anesthetic, consisting in a mixture of Zoletil (800 mg/kg) and Rompun 20 (20 mg/kg) administered intraperitoneally. To assess the expression of the immediate early gene protein c-Fos, euthanasia was performed 90 min after the end of the test. Once deep anesthesia was confirmed, the thoracic cavity was opened and animals were transcardially perfused with 0.1M phosphate-buffered saline (PBS) followed by 4% paraformaldehyde in PBS. Brains were then rapidly removed and post-fixed in 4% paraformaldehyde for 24 h. Subsequently, brains were transferred to 20% sucrose in 0.1 M PBS for 24 h and then to 30% sucrose in 0.1 M PBS until further histochemical analyses.

Coronal brain sections (35 µm) were cut using a cryostat (Leica CM1860 UV, Leica Biosystems, Germany). After sectioning, slices were stored in PBS containing 5% sucrose and 0.01% sodium azide at 4°C until immunohistochemical analysis. On day 1, sections were washed three times in PBST (0.1M PBS and TritonX) for 10 min each and then incubated for 2 h in blocking solution (0.1 M PBS, 5% normal goat serum). Sections were then incubated for 24 h with rabbit anti-c-Fos primary antibody (1:1000) diluted in a solution containing 0.1 M PBS, Triton X-100, bovine serum albumin (BSA), sodium azide, and sodium EDTA. On day 2, sections were washed three times in PBST for 10 min each and incubated for 2 h with the secondary antibody solution containing anti-rabbit Alexa Fluor 488 (1:500), diluted in the same vehicle used for the primary antibody solution. Sections were then washed three times in 0.1 M PBS for 10 min each. Finally, sections were mounted onto glass slides and left to dry overnight before coverslipping with Fluoromount-G containing DAPI (Invitrogen, Thermo Fisher Scientific). For c-Fos quantification, 3 brain sections containing the vHPC region were selected, corresponding to the plates 73, 74, 75 of the rat brain atlas (Paxinos and Watson, 2013), and the average c-Fos positive cells was obtained for each animal. Images were taken through a software (Capture 2.0) associated with an epifluorescence microscope (Olympus U-RFL-T Nikon 801), using 20x objective. c-Fos positive cells (GFP marker) were manually counted by an expert operator blind to experimental conditions, using the software Fiji (Schindelin et al., 2012). The same software was used to check the co-localization of GFP and DAPI.

### 2.4 Statistical analysis

Statistical analysis was performed using GraphPad Prism 9 (GraphPad Software, Boston, MA, USA). During training, object exploration time was analyzed to exclude spontaneous side or object preference before testing. Animals were considered to display a side preference when more than 60% of the total object exploration time was directed toward one of the two objects. In this case, the position of the displaced object was counterbalanced across animals and experimental groups. A two-way repeated-measures ANOVA was used to analyze object exploration time during OR and OL, with Odor exposure (NO, OD) as a between-subject factor and Object location (old, novel) as the a within-subject repeated factor. For the analysis of sex differences in memory performance and in c-Fos expression, %DI and c-Fos density were compared between males and females using two-way ANOVA, with Sex (male, female) and Odor exposure (NO, OD) as between-subject factors. When the ANOVA revealed significant main effects or interactions, post-hoc multiple comparisons were performed using Sidak’s test.

An unpaired t-test was used to compare c-Fos expression and %DI values between the NO and OD groups on OR or OL tests. One-sample t-test against zero was used to determine whether each group showed significant preference for the novel object or the novel object location. Before statistical analysis, data were screened for outliers using the ROUT method with Q = 1%. Normality was assessed using the Shapiro–Wilk test. When data did not meet the assumption of normality, the corresponding non-parametric test was used. For ANOVA models, residual normality and homoscedasticity were assessed by inspecting Q–Q plots and residual-versus-fitted-value plots, respectively. The final number of animals included in each analysis is indicated in the figure legends. Statistical significance was set at P < 0.05.

## 3. Results

### 3.1 Forced hyperconsolidation enhances object location memory, but not object recognition, in male rats

To assess the effects of forced hyperconsolidation on memory performance in male rats, we first analyzed exploration time, expressed in seconds, of each object (old vs novel) in the OR task, and found no significant effect of odor exposure (F_(1,25)_ = 3.00, P = 0.096), object familiarity (F_(1,25)_ = 0.257, P = 0.617) nor a significant interaction between the two factors (F_(1,25)_ = 0.097, P = 0.758) (**Fig. 2A**). The DI, expressed as percentage, confirmed the absence of significant differences between the two groups (t_(25)_ = 0.464, P = 0.646 **Fig. 2B**). Similarly, one-sample t-tests performed on DI% did not reveal significant discrimination in either the NO group (t_(8)_ = 0.201, P = 0.846) or the OD group (t_(17)_ = 0.825, P = 0.421 **Fig. 2B**). Total exploration of the two objects was not affected by odor exposure (t_(25)_ = 1.732, P = 0.096 **Fig. 2C**).

**Fig. 2.**
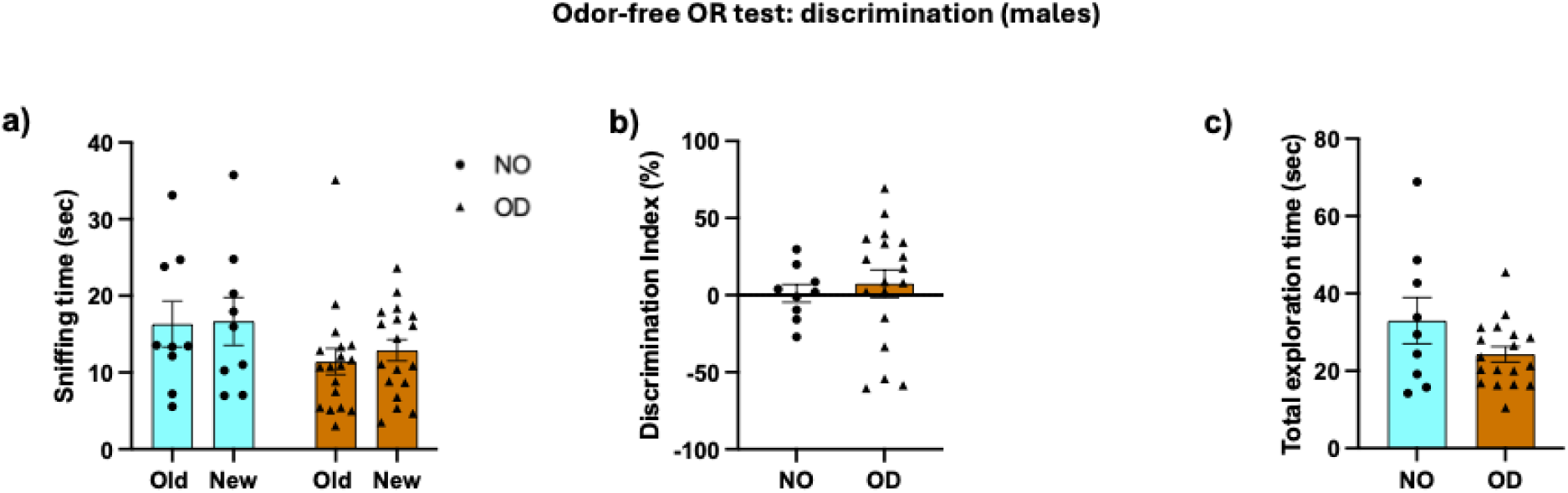
Effect of the forced hyperconsolidation protocol in male rats on object recognition memory. (a) In the OR test, the NO and OD groups did not differ in the time spent exploring the old or new object, and (b) they also did not differ in the discrimination index. (c) No difference in total object exploration time was observed between the NO and OD groups. Dots and triangles represent single animals, bars show mean and error bars represent SEM. N = 9 (NO), 18 (OD).

We also analyzed general locomotor activity in the arena during the test phase, and found no differences between the two groups in the number of horizontal activity (t_(25)_ = 0.171, P = 0.865), measured as crossings (**Fig. 3A**), or in the number (t_(25)_ = 1.24, P = 0.227) and duration (t_(25)_ = 2.043, P = 0.052) of vertical activity, measured as rearing and wall rearing (**Fig. 3B-C**)

**Fig. 3.**
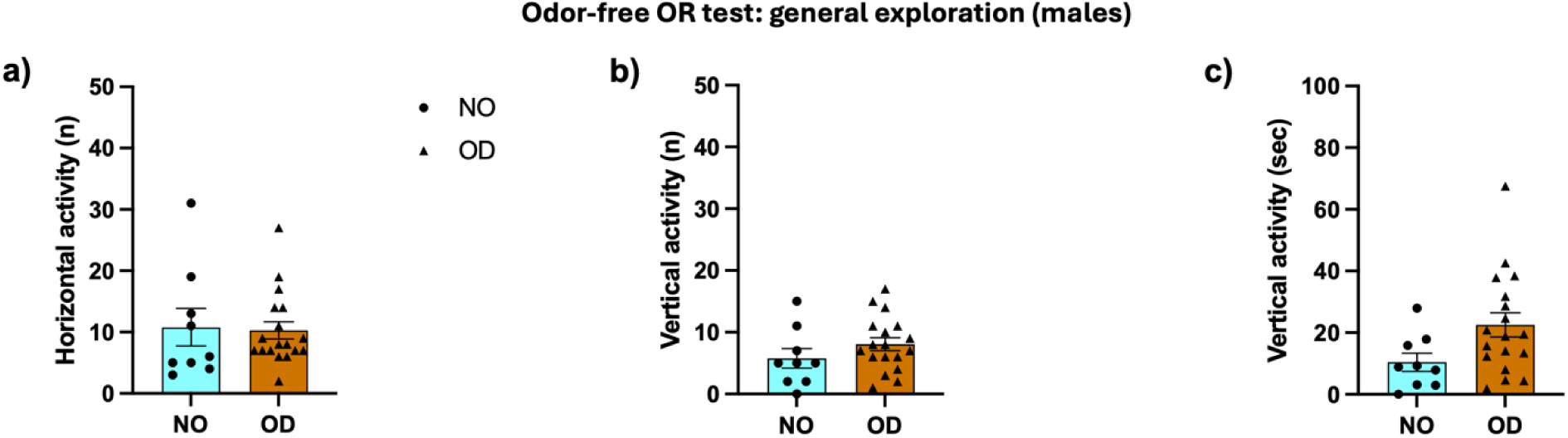
General exploratory activity in male rats tested in the object recognition paradigm. (a) The NO and OD groups did not differ in the horizontal activity, measured as the number of crossings in the experimental arena. No difference was found (b) in the number nor (c) in the duration of vertical activity, measured as rearings and wall rearings Dots and triangles represents single animals, bars show mean and error bars represent SEM. N = 9 (NO), 18 (OD).

In contrast, in the OL task, we found a significant effect of object location (F_(1,37)_ = 6.137, P = 0.018) and a significant odor exposure × location interaction (F_(1,37)_ = 14.16, P = 0.0006), but no main effect of odor exposure (F_(1,37)_ = 0.005, P = 0.945). Sidak’s post-hoc comparisons revealed that OD rats spent more time exploring the displaced location, relative to the old one (P < 0.0001), while no difference was found in the NO group (P = 0.637 **Fig. 4A**). Consistently, the DI% showed a significant difference between NO and OD rats (t_(37)_ = 3.34, P = 0.002 **Fig. 4B**). One-sample t-tests against casual discrimination further revealed significant discrimination in the OD group (t_(21)_ = 6.710, P < 0.0001), but not in the NO group (t_(16)_ = 0.522, P = 0.609 **Fig. 4B**). Total exploration of the two objects was not affected by odor exposure (t_(37)_ = 0.069, P = 0.945 **Fig. 4C**). No differences between the two groups were found in the horizontal activity (t_(37)_ = 0.212, P = 0.827 **Fig. 5A**), measured as the number of crossings, or in the number (t_(37)_ = 0.595, P = 0.555 **Fig. 5B**) and duration (t_(37)_ = 0.772, df = 37, P = 0.445 **Fig. 5C**) of vertical activity, measured as rearing and wall rearing.

**Fig. 4.**
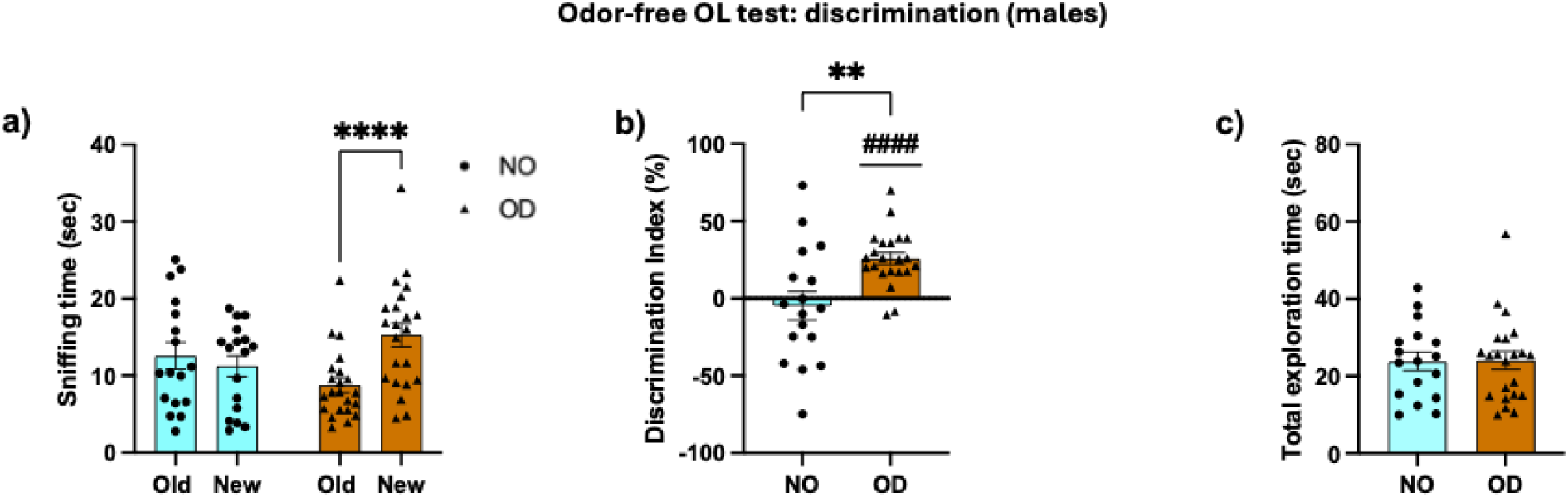
Effect of the forced hyperconsolidation protocol in male rats on object location memory. (a) In the OL test, only the OD group explored the object in the new position significantly more than the object in the old position, and (b) the OD group showed a higher discrimination index than the NO group, with values significantly greater than chance level. (c) No difference in total object exploration time was observed between the NO and OD groups. Dots and triangles represent single animals, bars show mean and error bars represent SEM. **, ****p < 0.01, 0.0001Unpaired t-test; ## p < 0.01 One-sample t-test. N = 17 (NO), 22 (OD).

**Fig. 5.**
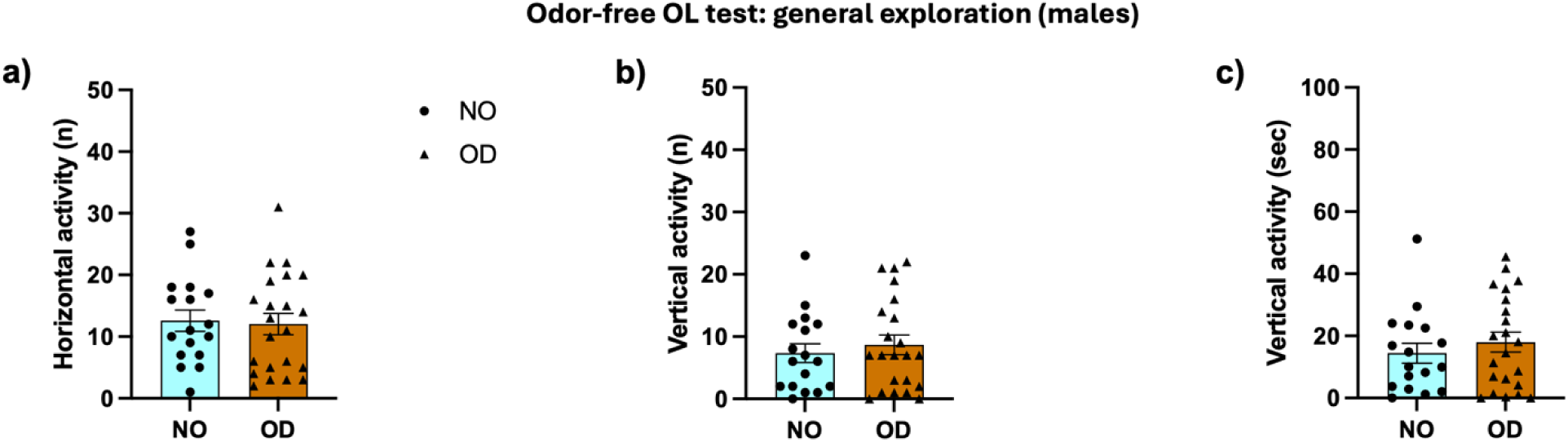
Exploratory and locomotor activity in male rats tested in the object location paradigm. (a) The NO and OD groups did not differ in the horizontal activity, measured as the number of crossings in the experimental arena. No difference was found (b) in the number nor (c) in the duration of vertical activity, measured as rearings and wall rearings. Dots and triangles represent single animals, bars show mean and error bars represent SEM. N = 17 (NO), 22 (OD).

### 3.2 Forced hyperconsolidation enhances object location memory, but not object recognition, in female rats

Similarly to males, forced hyperconsolidation had no effect in the discrimination of a new object. As such, in the OR test, we found no significant effect of odor exposure (F_(1,16)_ = 1.184, P = 0.293), object familiarity (F_(1,16)_ = 2.143, P = 0.163), nor significant interaction (F_(1,16)_ = 0.034, P = 0.855 **Fig. 6A**). The absence of significant differences between the two groups was confirmed by comparison of the %DI (t_(16)_ = 0.497, P = 0.625) (**Fig. 6B**). Similarly, one-sample t-tests performed on DI% did not reveal significant discrimination in either the NO group (t_(6)_ = 1.308, P = 0.239) or the OD group (t_(10)_ = 0.652, P = 0.529) (**Fig. 6B**). Total exploration of the two objects was not affected by odor exposure (t_(16)_ = 1.088, P = 0.293 **Fig. 6C**). The Unpaired t-tests revealed no differences between the two groups in the horizontal activity (t_(16)_ = 1.209, P = 0.244 **Fig. 7A**), measured as crossings, or in the number (t_(16)_ = 0.453, P = 0.657 **Fig. 7B**) and duration (t_(16)_ = 1.467, P = 0.162 **Fig. 7C**) of vertical activity, measured as rearing and wall rearing.

**Fig. 6.**
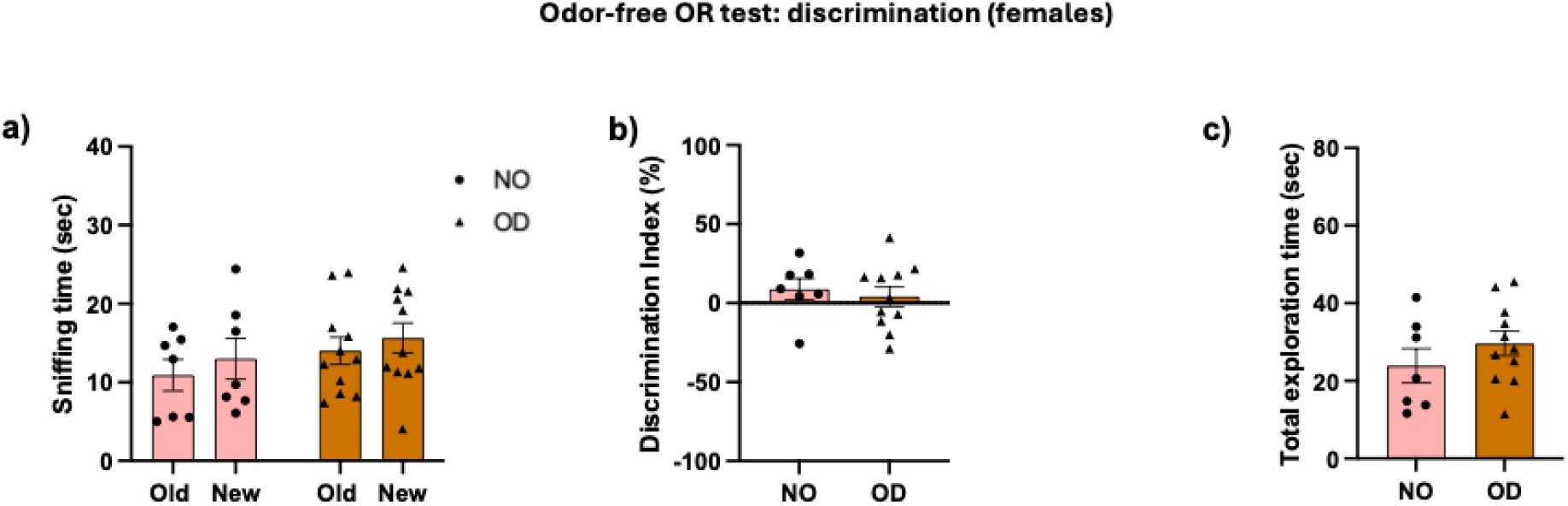
Effect of the forced hyperconsolidation protocol in female rats on object recognition memory. (a) In the OR test, the NO and OD groups did not differ in the time spent exploring the old or new object, and (b) they also did not differ in the discrimination index. (c) No differences in total object exploration time were observed between the NO and OD groups. Dots and triangles represent single animals, bars show mean and error bars represent SEM. N = 7 (NO), 11 (OD).

**Fig. 7.**
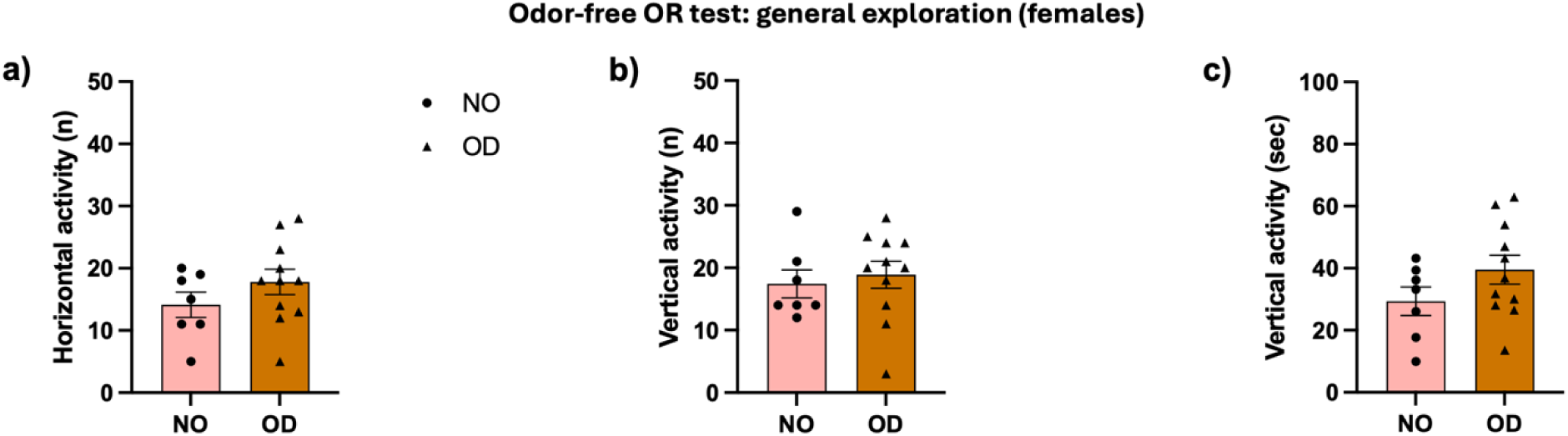
Exploratory and locomotor activity in female rats tested in the object recognition paradigm. (a) The NO and OD groups did not differ in the horizontal activity, measured as the number of crossings in the experimental arena. No difference was found (b) in the number nor (c) in the duration of vertical activity, measured as rearings and wall rearings. Dots and triangles represent single animals, bars show mean and error bars represent SEM. N = 7 (NO), 11 (OD).

In the OL test we found a significant effect of object location (F_(1,37)_ = 8.132, P = 0.007) and a significant odor exposure × location interaction (F_(1,37)_ = 6.778, P = 0.013), while no main effect of odor exposure was detected (F_(1,37)_ = 0.258, P = 0.615). Sidak’s post-hoc comparisons showed that female rats exposed to the odor spent more time exploring the novel location, compared to the old location (P = 0.0006), while no difference in location preference was found in the NO group (P = 0.982 **Fig. 8A**). The DI% was also significantly different between NO and OD females (t_(37)_ = 2.823, P = 0.008 **Fig. 8B**). By comparing each group against no discrimination, we found significant discrimination in OD rats (t_(20)_ = 3.275, P = 0.003), but not in NO rats (t_(17)_ = 0.307, P = 0.763 **Fig. 8B**). Total exploration of the two objects was not affected by odor exposure (t_(37)_ =0.185, P = 0.854 **Fig. 8C**). With respect to the general exploratory activity during the OL test, the two groups did not differ in the horizontal activity (t_(37)_ = 0.028, P = 0.978 **Fig. 9A**), measured as crossings, or in the number (t_(37)_ = 1.780, P = 0.083 **Fig. 9B**) and duration (t_(37)_ = 0.526, P = 0.602 **Fig. 9C**) of vertical activity, measured as rearing and wall rearing.

**Fig. 8.**
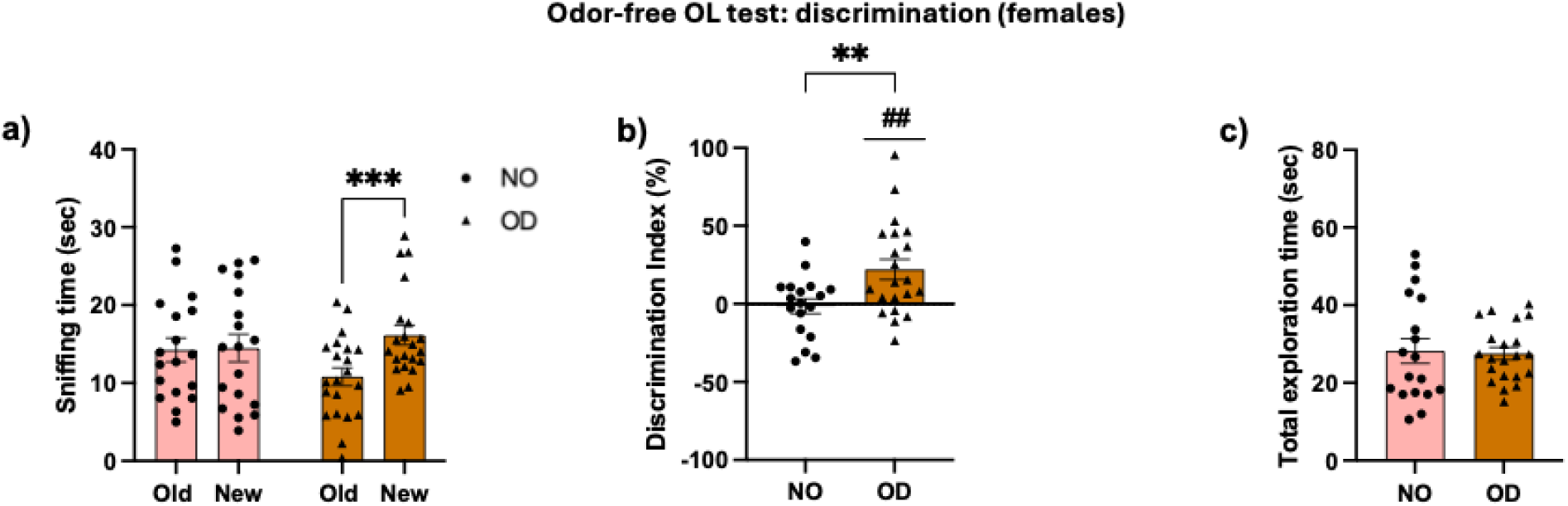
Effect of the forced hyperconsolidation protocol in female rats on object location memory. (a) In the OL test, only the OD group explored the object in the new position significantly more than the object in the old position, and (b) the OD group showed a higher discrimination index than the NO group, with values significantly greater than 0. (c) No difference in total object exploration time was observed between the NO and OD groups. Dots and triangles represent single animals, bars show mean and error bars represent SEM. **, *** p < 0.01, 0.001 Unpaired t-test; ## p < 0.01 One Sample t-test. N = 18 (NO), 21 (OD).

**Fig. 9.**
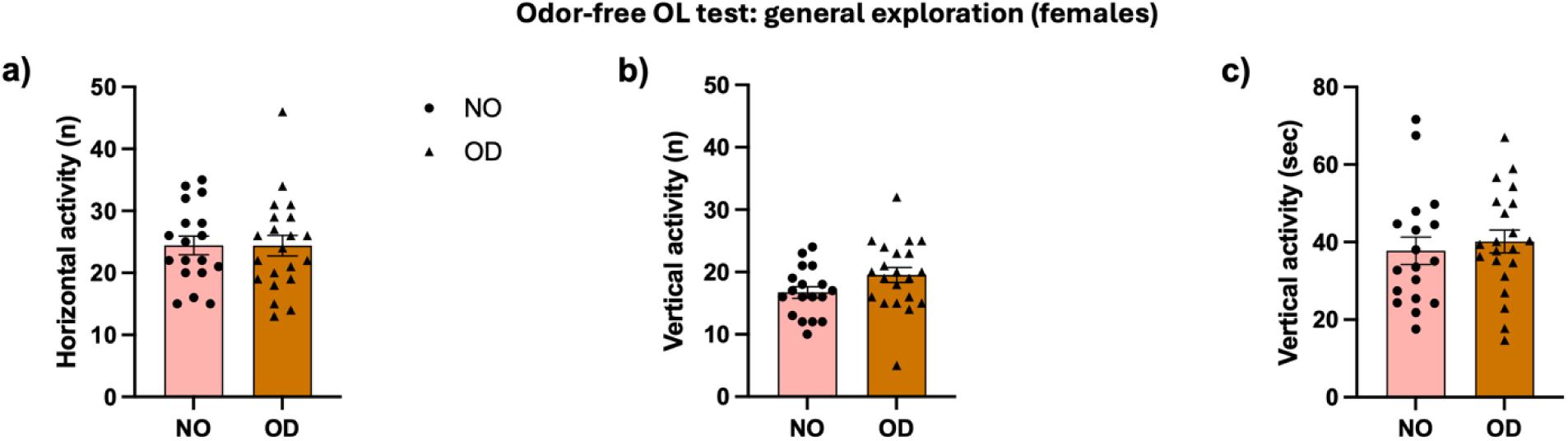
Exploratory and locomotor activity in female rats tested in the object location paradigm. (a) The NO and OD groups did not differ in the horizontal activity, measured as the number of crossings in the experimental arena. No difference was found (b) in the number nor (c) in the duration of vertical activity, measured as rearings and wall rearings. Dots and triangles represent single animals, bars show mean and error bars represent SEM. N = 18 (NO), 21 (OD).

### 3.3 Forced hyperconsolidation increases c-Fos expression in the vHPC of male and female rats during object location task

The density of Fos protein in the ventral hippocampus (vHPC), expressed as the number of c-Fos+ cells per mm², was quantified and compared in OD vs NO male rats, demonstrating a significant difference between the NO and OD (t_(9)_ = 2.876, P = 0.018 **Fig. 10G**. Similarly, in females, Fos expression was augmented in the vHPC of odor-exposed rats (t_(15)_ = 3.223, P = 0.006 **Fig. 10H**).

**Fig. 10.**
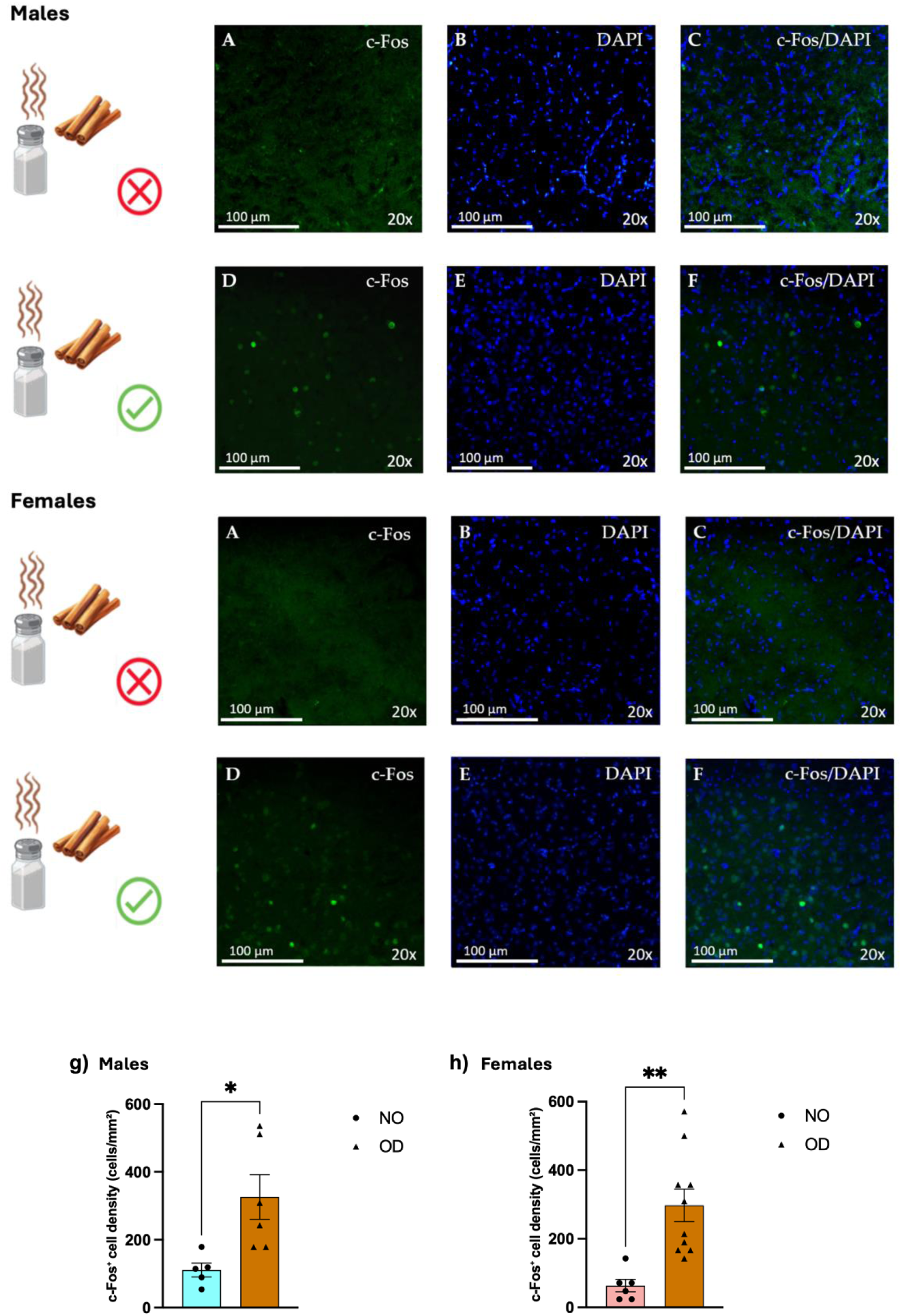
Object location task-induced c-Fos expression in the ventral hippocampus of odor-exposed and non-exposed rats. Representative fluorescence microscopy images of c-Fos expression in the vHPC of odor-re-exposed (OD) and non-odor-exposed (NO) male and female rats. Images were acquired at 20× magnification. (A, D) c-Fos immunoreactivity; (B, E) DAPI nuclear staining; (C, F) merged channels. Effect of forced hyperconsolidation on neuronal activation in the vHPC during the OL test in male (G) and female (H) rats. (G, H) In both male and female rats, the OD group showed a significantly higher number of c-Fos⁺ cells than the NO group. Dots and triangles represent single animals, bars show mean and error bars represent SEM. *, **, p <0.05, 0.01 Unpaired t-test. Males N = 5 (NO), 6 (OD); females N = 6 (NO), 10 (OD).

Additional analyses were performed to specifically assess sex differences. DI% values obtained in the OR and OL tests were compared between males and females of the two groups (OD vs NO). In the OR, the analysis revealed no significant effect of odor exposure (F_(1, 41)_ = 0.007, P = 0.933), sex (F_(1, 41)_ = 0.056, P = 0.814), nor odor exposure × sex significant interaction (F_(1, 41)_ = 0.371, P = 0.546 **Fig. 11A**). In the OL, consistently with what obtained with previous analysis, we found a significant main effect of odor exposure (F_(1,84)_ = 18.07, P < 0.0001), but no significant effect of sex (F_(1,84)_ = 0.027, P = 0.87) or significant odor exposure × sex interaction (F_(1,84)_ = 0.0271, P = 0.8697). Sidak’s post-hoc comparisons highlighted the enhanced discrimination by the OD group, both in males (P = 0.001) and in females (P = 0.042 **Fig. 11B**).

**Fig. 11.**
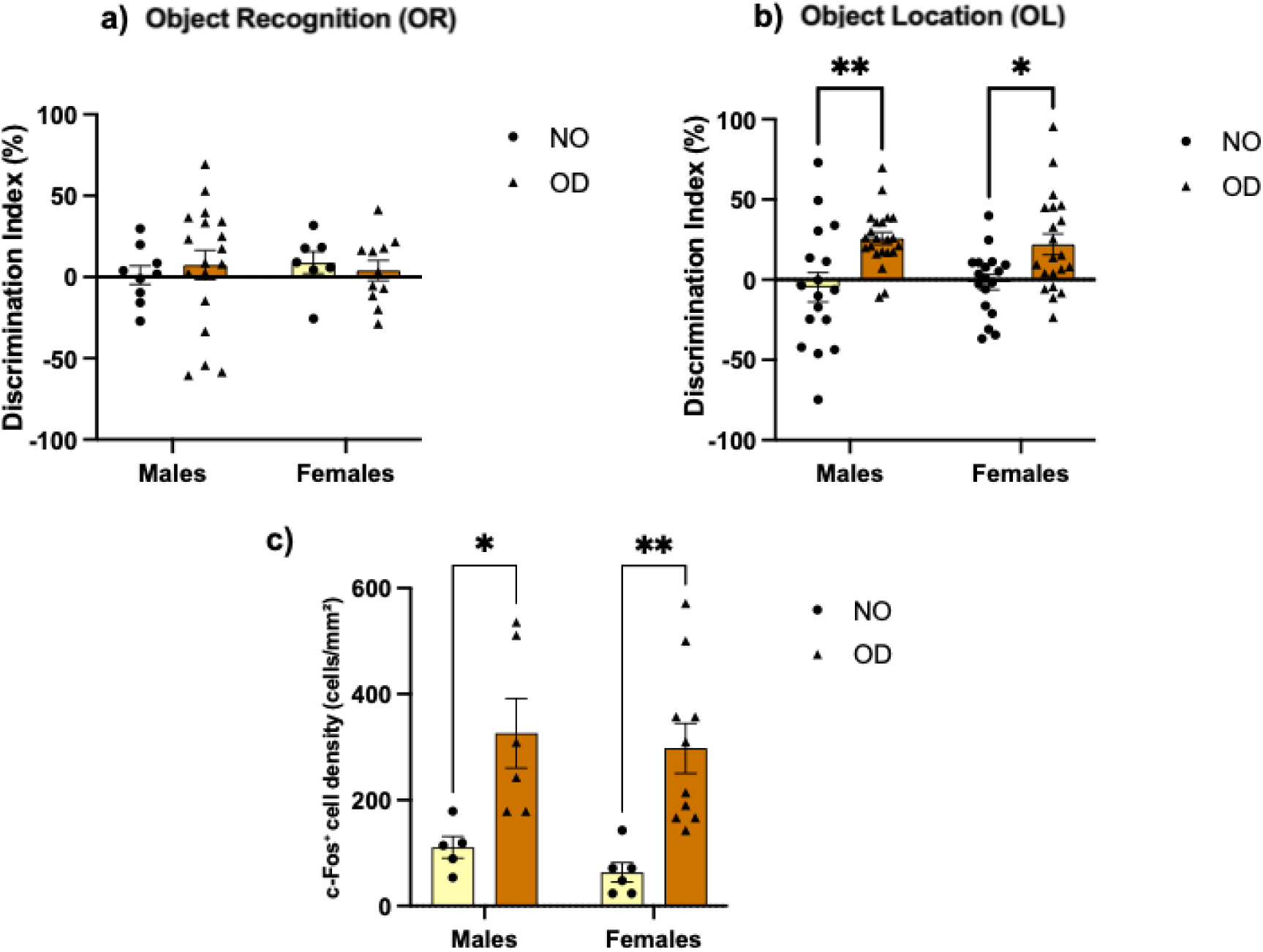
Comparison of the discrimination index between male and female rats in the NO and OD groups during the object recognition and object location paradigms. (a) No sex differences in the discrimination index were observed in the OR test. (b) In the OL test, no sex differences in the discrimination index were found. Both male and female rats in the OD group successfully discriminated the object in the novel position. (c) In the OL test, the increase in neuronal activation observed in the OD group compared with the NO group, expressed as c-Fos⁺ cell density, was comparable between males and females. Dots and triangles represent single animals, bars show mean and error bars represent SEM. *, ** p <0.05, 0.01 Sidak’s post-hoc multiple comparisons. OR: males N = 27, females N = 18; OL: males N = 39, females N = 39; c-Fos: males N = 11, females N = 16

The same analysis was also performed on c-Fos expression. Two-way ANOVA revealed a statistically significant effect of odor exposure (F_(1,23)_ = 21.13, P = 0.0001) but not of sex (F_(1,23)_ = 0.601, P = 0.446), and no significant odor exposure × sex interaction (F_(1,23)_ = 0.037, P = 0.848). Sidak’s post-hoc analysis confirmed the increased c-Fos expression in the OD group of both sexes (males, P = 0.016; females, P = 0.002 **Fig. 11C**).

## 4. Discussion

Pathological alterations of episodic memory are not limited to memory impairments, such as those observed in Alzheimer’s disease and other neurodegenerative disorders (Khan et al., 2026), but may also involve memory hyperfunction (Santangelo et al., 2025). This is the case in disorders characterized by a ruminative component, such as PTSD and depression, in which some memories may become excessively persistent, intrusive, and difficult to extinguish (Berardi et al., 2014; Morena et al., 2018). However, animal models specifically designed to mimic rumination-induced memory alterations are still lacking. Accordingly, the aim of the present study was therefore to develop a behavioral paradigm able to reproduce the hyperconsolidation of selected episodic-like memories in both male and female rats, taking inspiration from the persistence of memory observed in individuals with HSAM. Moreover, since this study was intended to provide a tool to investigate the mechanisms underlying hypermemory-like phenomena, we also assessed neuronal activity, through c-Fos analysis, in the vHPC, a region strongly implicated in the retrieval of episodic and contextual memories (Howland et al., 2008; Komorowski et al., 2013; Veyrac et al., 2015).

The procedure established here to force hyperconsolidation consisted in re-exposing animals to the odor presented during the training phase of the OR or OL tasks. The choice of olfactory stimuli as retrieval cues deserves specific consideration. Among sensory modalities, olfaction holds a unique neuroanatomical position, conferring olfactory information with exceptional efficacy in triggering the reactivation of hippocampus-dependent contextual memory traces, a phenomenon well documented in humans as the “Proustian memory effect” (Aqrabawi and Kim, 2018; Baudry, 2020). In rodents, odor-induced memory reactivation has been shown to engage reconsolidation mechanisms, rendering the original memory trace susceptible to strengthening or updating (Strauch and Manahan-Vaughan, 2020). The repeated pairing of a distinctive odor with the learning context in our paradigm was therefore designed to exploit this reactivation-consolidation cycle, hypothesizing that each olfactory re-exposure would trigger a new consolidation event and progressively strengthen the memory trace. We hypothesized that reactivating the object memories through the previously associated odor would enhance retrieval performance in both tasks. Contrary to our hypothesis, however, memory retention was enhanced only in the OL task, as shown by the increased exploration of the displaced object and the higher DI% observed in odor-re-exposed animals compared with non-re-exposed controls.

The different outcomes obtained in the OR and OL tasks are interesting, as they suggest a selective effect of hyperconsolidation process, highlighting a task-specific pattern that might be neuroanatomically informative. Our findings support the idea that the hippocampus is primarily involved in spatiotemporal processing and contextual mapping. Although the dorsal hippocampus is believed to play a more direct role in spatial processing, being richer in place cells (Hughes, 1965; O’Keefe, 1976; Sinnamon et al., 1978), communication with the ventral portion (i.e., the hippocampal region in which we observed increased c-Fos expression in both male and female rats during the OL task) may facilitate the retrieval of this information through interactions with the prefrontal cortex (Eichenbaum, 2017; Fanselow and Dong, 2010; Bakoyiannis et al., 2023). This aligns with the core mechanism proposed for hypermemory, which involves altered functional connectivity between the prefrontal cortex and the hippocampus (Daviddi et al., 2022; Orwig et al., 2024; Parente et al., 2026; Santangelo et al., 2018, 2020, 2021).

From a neuroanatomical standpoint, this task-selectivity is further supported by the trajectory of olfactory projections: odor information reaches the hippocampus through a relatively direct route via the olfactory bulb, piriform cortex, and lateral entorhinal cortex, with comparatively weaker projections to the perirhinal cortex (Ergorul and Eichenbaum, 2004; Chapuis and Wilson, 2012). This asymmetry in connectivity may explain why repeated odor reactivation selectively strengthened hippocampus-dependent contextual memory without influencing perirhinal cortex-dependent object recognition (Li et al., 2017; Symanski et al, 2022). Indeed, in the OL task, the preference for the displaced object confirms the involvement of the hippocampus in the processing and retrieval of spatial information and orientation, unlike object recognition memory, which is more strongly associated with the perirhinal cortex (Danieli et al., 2023). In this context, the vHPC may play a key role in linking object information with contextual representations, possibly by enhancing signal transmission to the prefrontal cortex (Morici et al., 2022).

Importantly, we have shown that the effect of odor re-exposures on object exploration was not biased by altered locomotion or general not object-related exploratory activity, as shown by the analysis of both horizontal and vertical behaviors in both OR and OL tasks. Moreover, the total exploration time of the two objects did not change as a function of odor re-exposures, indicating that the forced hyperconsolidation protocol did not affect the general interest for the objects. The enhanced OL performance in the OD group did not reflect a single unidirectional change in object exploration; rather, it emerged from a combined pattern of reduced exploration of the familiar location and increased exploration of the displaced object. This suggests that repeated odor-cued reactivations produced a genuine strengthening of the spatial memory trace, enabling more efficient recognition of the changed configuration. Through the c-Fos analysis, we aimed to investigate the involvement of the vHPC in the retrieval of hyperconsolidated contextual information related to an episodic-like memory. c-Fos, an immediate early gene whose protein product is transiently expressed following neuronal activation, represents a well-established and minimally invasive tool for mapping activity-dependent neuronal recruitment at the circuit level (Polanczyk et al., 2026). Its use here serves as a neurobiological readout complementary to the behavioral measures, allowing verification that the paradigm engages the expected neural substrates and not merely produces a non-specific motor or motivational effect. This analysis revealed greater activation of the vHPC in odor-re-exposed animals compared with non-exposed controls in both males and females, thereby supporting the behavioral findings and providing neurobiological validation of the hyperconsolidation protocol. The convergence of behavioral enhancement and increased vHPC c-Fos expression confirms that the paradigm selectively recruits hippocampal circuitry during the reactivation window, consistent with the role of the vHPC in integrating contextual and emotional dimensions of episodic-like memory (Veyrac et al., 2015; Auguste et al., 2023).

The replication of this neural signature across sexes further strengthens the methodological reliability of the approach. Although HSAM has been reported in both males and females, available evidence suggests that more males than females have been described with this condition. For instance, LePort et al. (2016) reported a cohort of 30 HSAM individuals, including 24 males. However, current evidence does not yet support firm conclusions about sex-specific prevalence or mechanisms, as exact sex distributions remain uncertain due to the rarity of HSAM, the limited number of identified individuals, small sample sizes, and the lack of clarity regarding possible participant overlap across studies (Talbot et al., 2024). In our experimental studies, no significant sex differences were observed, leaving open the possibility that forced hyperconsolidation relies on largely shared neurobiological mechanisms in males and females.

The present model demonstrates face validity in that it recapitulates, at the phenotypic level, the core feature of HSAM: a selective and disproportionate enhancement of hippocampus-dependent contextual memory, as reflected by improved object location performance in the absence of a corresponding change in object recognition. This selectivity mirrors the profile reported in HSAM individuals, who exhibit superior autobiographical and contextual memory without a generalized enhancement of all memory domains (LePort et al., 2012; McGaugh and LePort, 2014). Furthermore, the use of repeated olfactory reactivation as the induction mechanism parallels the spontaneous rumination cycles thought to underlie HSAM in humans. Together, these features position the model as a viable preclinical platform for investigating the neurobiological substrates of hypermemory and for testing interventions aimed at modulating pathological forms of memory persistence.

Several rodent models have been developed to study enhanced memory consolidation, most notably fear-based paradigms used in PTSD research, in which aversive conditioning or stress exposure produces persistent and intrusive fear memories mediated by amygdala-hippocampal hyperactivation (Maren et al., 2013; Berardi et al., 2014). While these models have substantially advanced our understanding of maladaptive memory persistence, they are inherently dependent on aversive experience, limiting their applicability to the study of non-fear-based hypermemory. Similarly, pharmacological facilitation protocols — such as post-training glucocorticoid administration or nootropic treatment — enhance memory consolidation broadly but through systemic neurochemical manipulations that lack mechanistic specificity and translational naturalism (Roozendaal et al., 2009; Campolongo et al., 2009; Atsak et al., 2015). The present paradigm differs from both approaches in two key respects: it induces selective enhancement of hippocampus-dependent contextual memory without aversive components, and it does so through a naturalistic, experience-dependent mechanism (olfactory reactivation) that recapitulates the rumination-like induction process discussed above. This positions the model in a largely unoccupied niche between fear-memory enhancement and generic cognitive facilitation, specifically suited to investigate non-aversive forms of memory superiority.

## 5. Conclusions

Overall, rather than representing a direct preclinical reproduction of HSAM in the strict sense, this paradigm should be viewed as a behaviorally grounded, non-invasive experimental tool to investigate the mechanisms underlying enhanced and persistent memory consolidation in rodents. Unlike pharmacological or genetic approaches to memory enhancement, which involve constitutive manipulations with broad neurochemical consequences, the proposed protocol exploits a biologically grounded, experience-driven mechanism that is both plausible and experimentally reproducible. Indeed, it should be acknowledged that autobiographical and subjective components that characterize human episodic memory cannot be fully assessed in animals. Nevertheless, this does not reduce the value of the model, but rather defines its specific field of application. By promoting the strengthening and persistence of episodic-like memory traces through olfactory reactivation, a mechanism supported by convergent behavioral and neurobiological evidence across sexes, the paradigm provides a robust and versatile framework to explore processes that resemble maladaptive memory persistence and rumination-like phenomena, allowing the identification of underlying neurobiological mechanisms and potential targets for intervention in conditions such as PTSD, OCD, and rumination-related disorders (Donaldson and Hen, 2015; Schmitz et al., 2017; Morena et al., 2018).

## CRediT authorship contribution statement

**Elvira Conforti:** Writing – original draft, Writing – review & editing, Methodology, Investigation, Formal analysis, Data curation. **Giulia Chiacchierini:** Writing – review & editing, Methodology, Investigation, Formal analysis, Data curation, funding acquisition. **Alberto Marchetti:** Investigation, Formal analysis, Data curation. **Jacopo Maria Lisiola:** Methodology, Investigation, Formal analysis. **Valerio Santangelo:** Writing – review & editing, Supervision, Formal analysis, Funding acquisition, Data curation, Conceptualization. **Patrizia Campolongo:** Writing – review & editing, Validation, Supervision, Resources, Funding acquisition, Data curation, Conceptualization.

